# A stem cell aging clock links biological age deviation to clinical outcome in acute myeloid leukemia

**DOI:** 10.64898/2026.02.12.703707

**Authors:** Hongcheng Chen, Pengwei Dong, Jianing Xu, Guanlin Wang

**Affiliations:** Shanghai Key Laboratory of Metabolic Remodelling and Health, Institute of Metabolism and Integrative Biology, School of Basic Medical Sciences, Centre for Evolutionary Biology, Fudan University, Shanghai, 200438, China

**Author notes:** Corresponding author: Guanlin Wang.

## Abstract

Aging of hematopoietic stem and progenitor cells (HSPCs) impairs regenerative capacity and predisposes to hematological diseases. Here, we constructed a comprehensive single-cell transcriptomic atlas comprising 186,123 CD34^+^ HSPCs spanning early prenatal development (6 post-conception weeks) to late adulthood (74 years). We identified two conserved core molecular programs (MPs) of inflammaging and RNA splicing / protein homeostasis. Leveraging these programs, we developed a machine learning-based stem cell aging clock from 84 donors. Applying this clock to acute myeloid leukemia (AML), we define Transcriptional Age Deviation (TAD), a novel metric of biological age divergence. We found that a biologically “younger” state (low TAD), reflecting oncofetal reprogramming, is a powerful independent predictor of poor survival in two large AML cohorts. Low TAD was associated with high-risk genetics and therapy resistance, and critically, it re-stratified patient outcomes within each ELN 2022 risk category. Our work establishes a quantitative link between stem cell aging biology and AML prognosis, offering a robust tool to refine clinical risk assessment.

## Introduction

Hematopoietic stem and progenitor cells (HSPCs) serve as the foundation of lifelong hematopoiesis, sustaining hematopoietic homeostasis and immune competence throughout the human lifespan^1–4^. However, aging imposes a progressive decline in HSPC function, leading to reduced regenerative capacity, lineage skewing toward myeloid differentiation, accumulation of DNA damage and increased susceptibility to hematological disorders such as acute leukemia^5,6^. Notably, the aged hematopoietic system contributes to systemic aging across multiple organs, as demonstrated by heterochronic parabiosis experiments showing that young blood can rejuvenate aged tissues^7^, highlighting the central role of HSPCs in organismal aging. Multiple mechanistic studies in mouse have identified several key drivers and pathways involved in HSPC aging, including increased inflammatory exposure (inflammaging) and disturbance of RNA splicing^8–13^. However, the extent to which these mechanisms operate in human HSPC aging remains to be fully elucidated.

Understanding the transcriptional programs governing both physiological aging and pathological transformation in HSPCs is crucial for identifying molecular vulnerabilities underlying age-related hematopoietic decline and malignant progression^14^. Studying human HSPC aging remains challenging due to the inherent heterogeneity of HSPCs and limited access to human samples across the lifespan. Traditional bulk RNA sequencing approaches fail to capture the complexity of individual HSPC gene expression profiles, missing subtle but biologically meaningful age-related transcriptional shifts. The advent of single-cell RNA sequencing (scRNA-seq) technologies has revolutionized this field by enabling high-resolution profiling of HSPC subpopulations across human development and aging, revealing a higher proportion of lymphoid-biased cells during early development, followed by a progressive shift toward myeloid-predominant hematopoiesis in the bone marrow niche during aging^15–19^. Nevertheless, the regulatory programs driving age-associated HSPC heterogeneity and functions remain incompletely understood.

Recent studies have introduced ‘aging clocks’ in various tissues^20–23^, but it remains unclear whether a transcriptomics-based stem-cell-specific clock can be robustly applied across the human lifespan, from prenatal to elderly age. More importantly, while clinical risk stratification in acute myeloid leukemia (AML), such as the ELN 2022 guidelines, has improved, significant heterogeneity remains within each risk group^24–27^. There is a critical need for novel, biologically-grounded biomarkers that can capture this heterogeneity and provide additional prognostic value. It is unknown whether a metric derived from the fundamental process of stem cell aging could fulfill this clinical need.

In this study, we constructed a comprehensive single-cell transcriptomic atlas of human CD34^+^ HSPCs spanning developmental stages from early prenatal development (6 PCW) to late adulthood (74 years) across 28 healthy donors. Molecular characterization revealed six conserved aging-associated molecular programs (MPs) representing distinct transcriptional signatures related to inflammatory signaling, HSC identity, transcription signaling, aerobic respiration, RNA splicing and proteostasis, and cell cycle regulation. Leveraging these programs, we developed a machine learning-based stem cell aging clock that robustly predicts donor chronological age from single-cell HSC transcriptomes and quantifies biological age from 84 donors. Applying this framework to AML, we defined transcriptional age deviation (TAD) as the age deviation of leukemic stem cells relative to a healthy baseline. TAD reveals disease-associated remodeling of stem cell aging states that extends beyond intratumoral heterogeneity, capturing clinically meaningful variation linked to genetic risk stratification and patient outcome. Collectively, our work provides a reference map of human HSPC aging, establishes a stem cell aging clock that enables quantitative integration of age-associated stem cell states at single-cell resolution, and provides a framework for uncovering aging-related dysregulation linked to disease outcome in hematological disorders.

## Results

### A single-cell transcriptomic atlas of HSPCs across human lifespan

To systematically characterize age-related dynamics in human hematopoiesis, we integrated six previously published datasets of CD34^+^ hematopoietic stem and progenitor cells (HSPCs), comprising a total of 200,665 cells from 28 healthy donors aged from post-conception week (PCW) 6 to 74 years (**Figure 1A-B**)^5,17,44–47^. To verify sample integrity, donor sex was confirmed based on expression of sex-chromosome markers *XIST* (female) and *UTY* (male) (**Supplemental Figure 1C**). After stringent quality control (**Supplemental Figure 1A-B, Supplemental Methods**), 186,123 high-quality cells were retained for downstream analyses (**Figure 1C-D, Supplemental Figure 1D-G, Supplemental Tables 1 and 2**). From this atlas, we isolated 59,723 high-confidence hematopoietic stem cells and multipotent progenitors (HSC/MPPs) based on unsupervised clustering and a validated gene signature score (**Figure 1E**)^17^. The primitive, undifferentiated state of this population was confirmed by differentiation potential (CytoTRACE2) and trajectory analyses (StaVIA, **Supplemental Figure 2A-C, Methods**)^31,32^.

**Figure 1.**
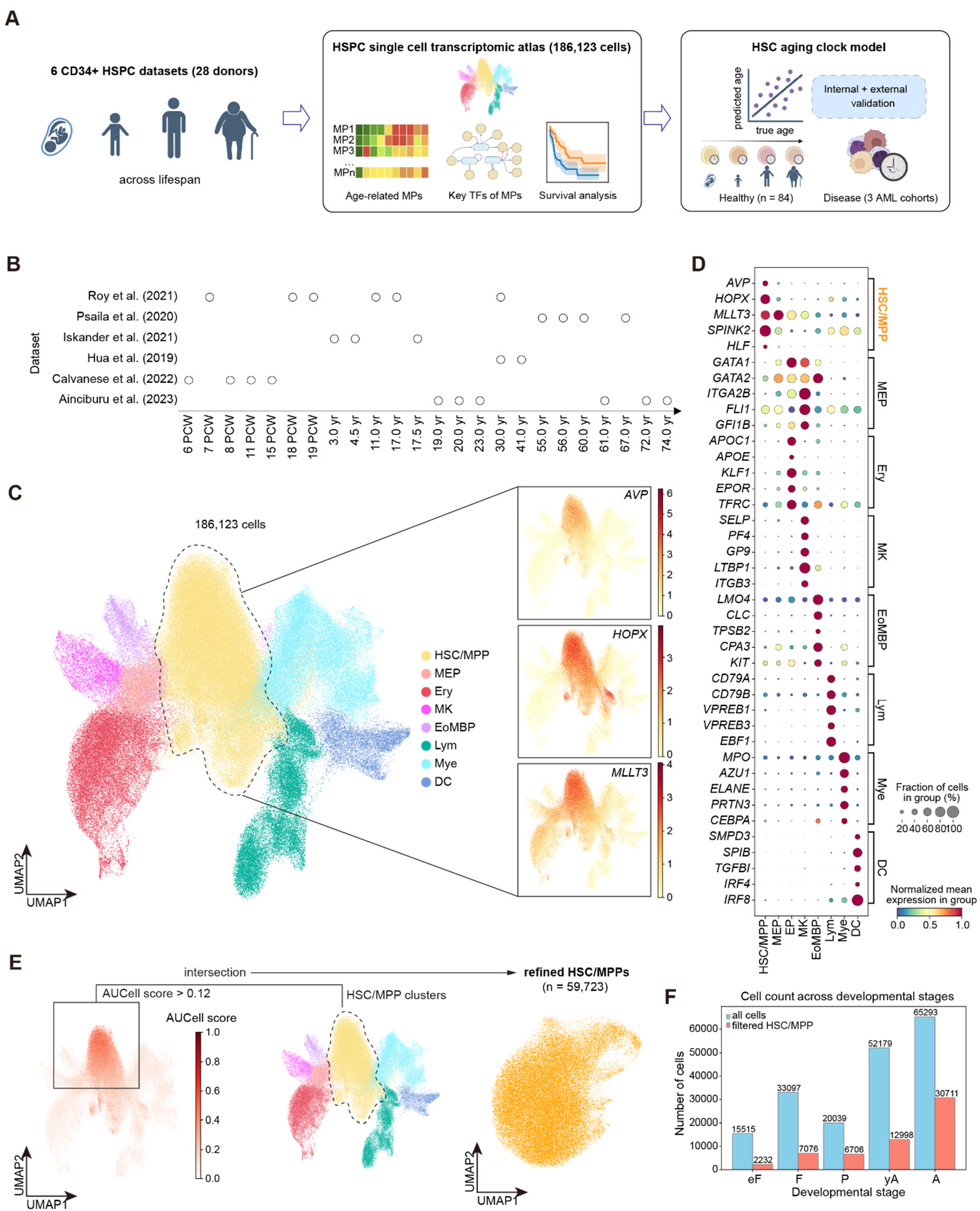
Construction of a human lifespan HSPC single-cell transcriptome atlas. (**A**) Schematic workflow of the study. (**B**) Sample sources for this study. PCW, post-conception week; yr, age. (**C**) UMAP dimensionality reduction plot of the human HSPC transcriptome atlas and highlighted expression of HSC marker genes (*AVP, HOPX, MLLT3*) on the UMAP projection. (**D)** Normalized expression of lineage-specific marker genes in HSPCs. (**E**) HSC/MPP filtration based on clustering and AUCell enrichment. Distribution of HSPC AUCell scores on UMAP projection (left), clustering result (middle), and UMAP projection of refined HSC/MPPs (right) were shown. HSC/MPP, hematopoietic stem cells and multipotent progenitors; MEP, megakaryocyte-erythroid progenitors; EP, erythroid progenitors; MK, megakaryocytes; EoMBP, eosinophil-basophil-mast progenitors; Lym, lymphoid progenitors; Mye, myeloid progenitors; DC, dendritic cells; eF, early fetal (PCW 6-12); F, fetal (PCW 13 to birth); P, pediatric (age ≤18 years); yA, young adult (18 < age ≤40 years); A, adult (age >40 years). Figure 1A were created using Biorender (www.biorender.com).

**Figure 2.**
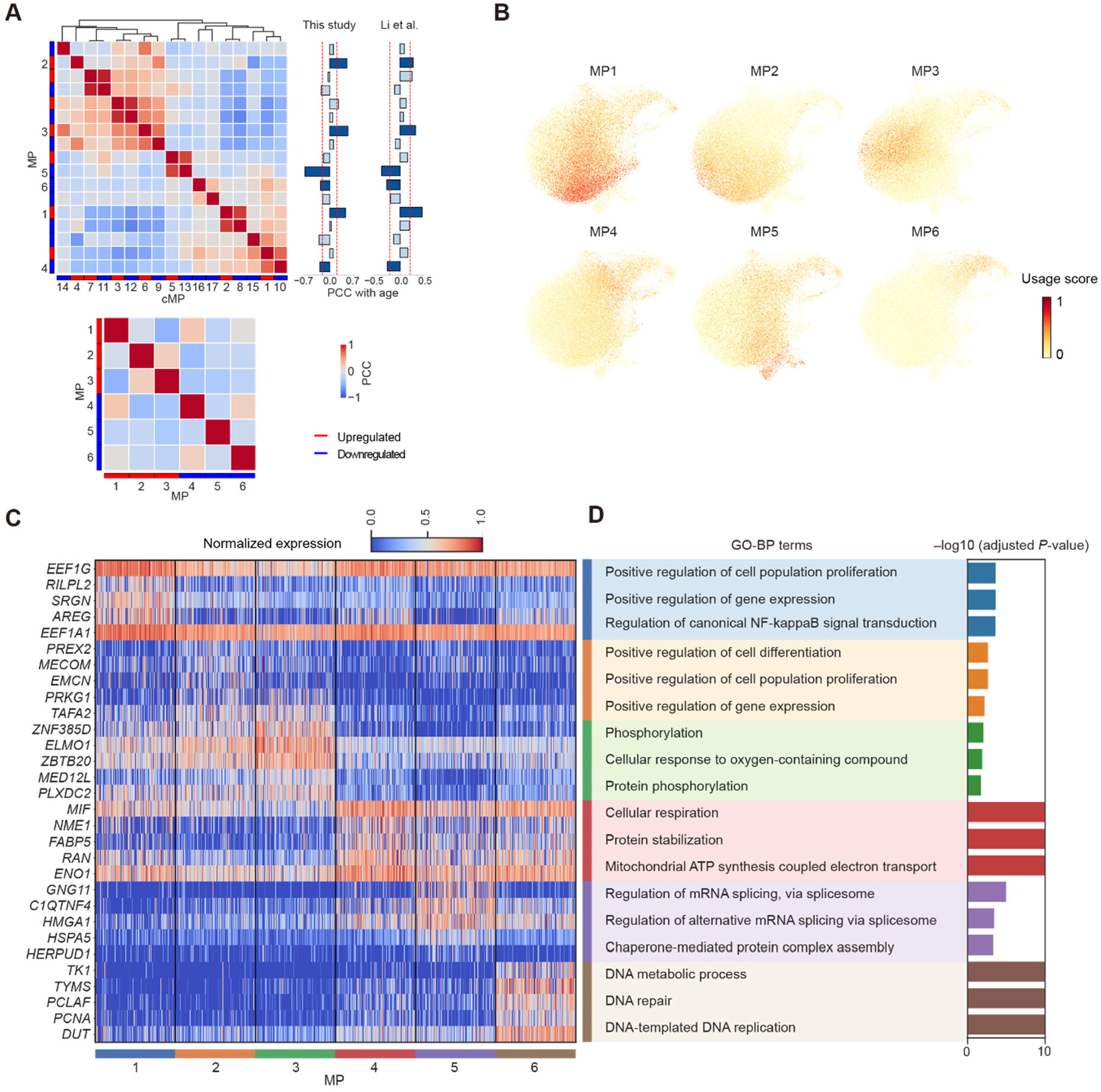
Age-related molecular programs in lifespan HSC/MPPs. (**A**) Correlation of 17 candidate molecular programs (cMPs, upper part) and 6 robust molecular programs (MPs, lower part). Correlation between cMP usage scores and age is shown on the right. 6 cMPs with Pearson’s Correlation Coefficient (PCC) > 0.20 and *P* < 0.05 in both discovery and validation cohorts were chosed. (**B**) Distribution of 6 MPs on UMAP embeddings. (**C** and **D**) Expression of top 5 genes from each MP on program-specific HSC/MPPs (**C**) and pathway enrichment results (**D**) of each MP. A cell was considered specific to the MP with the highest usage score. Pathway enrichment was performed using the GO Biological Process database. All pathways shown reached significance of adjusted *P* < 0.05.

### Six age-related molecular programs define HSC/MPP functional states

To delineate the age-related transcriptional reprogramming of the refined HSC/MPP population, we conducted UMAP projection, differential expression, and pySCENIC regulon analysis (**Supplemental Figure 2D-H, Supplemental Methods**)^30,48^. The analysis revealed a sharp age-dependent transcriptional and regulatory rewiring of HSC/MPP. Proliferative, MYC/E2F-dominated transcription dominates early fetal cells, whereas adult HSC/MPPs exhibit a quiescent, myeloid-primed state marked by heightened NF-κB inflammatory regulons (e.g. NFKB2, ERG), capturing the core transition from development to inflammaging in human hematopoiesis.

To delineate the latent transcriptional architecture of HSC/MPP aging beyond single gene alteration, we employed consensus non-negative matrix factorization (cNMF)^49^ to identify molecular programs (MPs) representing underlying functional states. We calculated six robust age-related MPs from our HSC/MPP discovery cohort and an independent validation cohort^16^ (**Figure 2A-B, Supplemental Figure 3A-B, Supplemental Table 4, Supplemental Methods**). Pathway analysis showed that the six MPs map to distinct functions, with MP1 (most significantly upregulated) reflecting pro-inflammatory NF-κB–driven “inflammaging,” MP5 (most significantly downregulated) characterizing RNA-splicing/proteostasis, and the remaining modules corresponding to HSC-identity, transcription-signal, mitochondrial respiration, and cell-cycle processes (**Figure 2C-D, Supplemental Table 5**).

**Figure 3.**
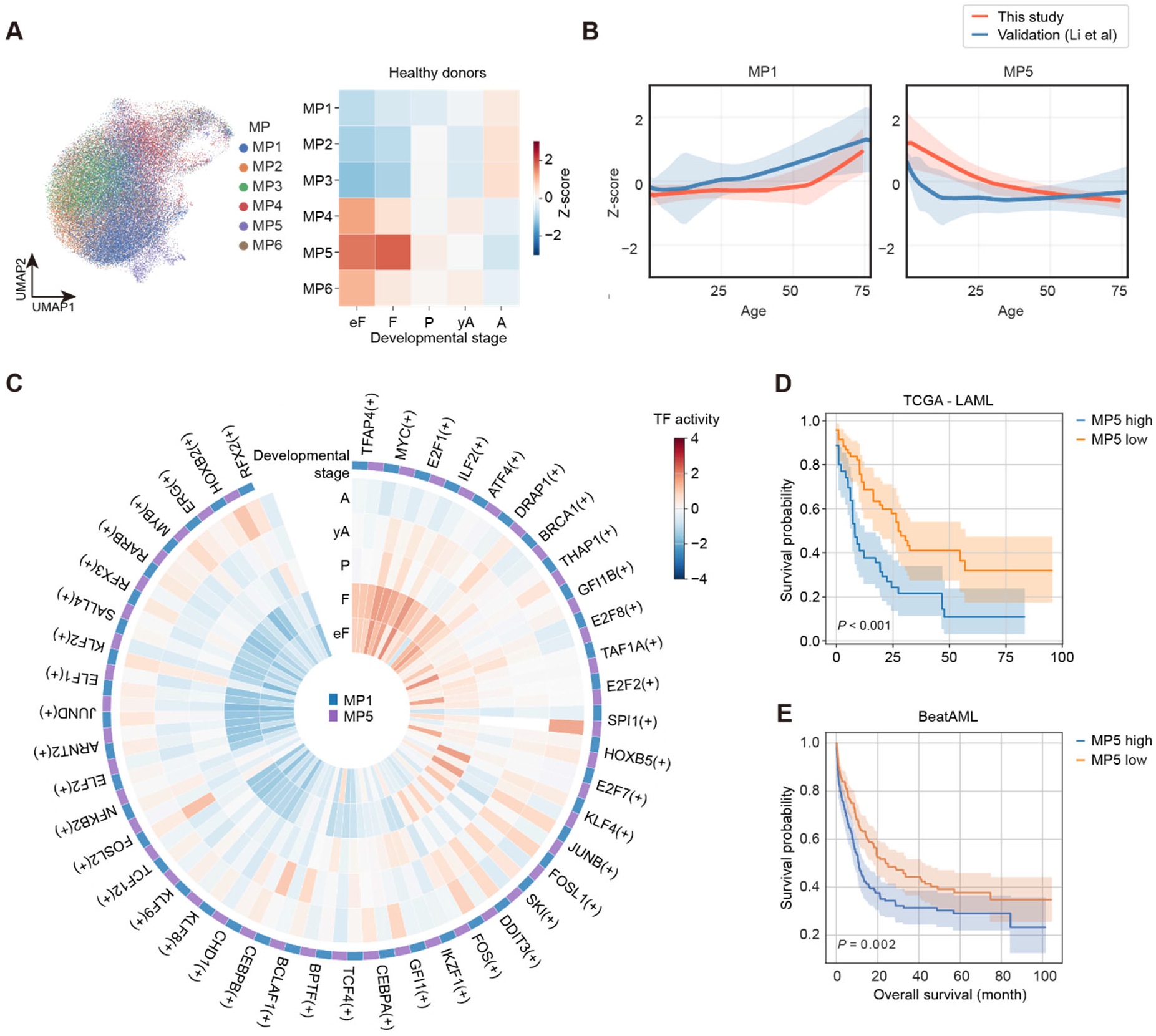
Characterization of molecular programs across developmental stages. (**A**) Distribution and activity of MPs across the five developmental stages. (**B**) Temporal dynamics of the two age-related MPs. (**C**) Activities of key transcription factors driving MP1 and MP5 across developmental stages. (**D** and **E**) Survival analysis results of MP5-related 18-gene set on TCGA-LAML (**D**) and BeatAML (**E**) cohorts.

### Age-associated dynamics of molecular programs in HSC/MPPs

To systematically explore age-associated changes in molecular programs in refined HSC/MPPs, we assessed the distribution and activity of MPs across five developmental stages and major hematopoietic lineages (**Figure 3A-B, Supplemental Figure 3C-D**). MP1-3 were predominantly enriched in adult ages, particularly within the uncommitted HSC/MPP and the myeloid compartment, suggesting progressive activation of myeloid-biased inflammaging pathways with age. In contrast, MP4-6 showed pronounced enrichment during early development (early fetal and fetal stages) but markedly decreased in postnatal and diminished in adult stages with modest variation across lineage contexts, indicating developmental-specific programs associated with aerobic respiration, transcription regulation, and proliferation that are attenuated after birth.

We next examined transcription factor (TF) activity within cells assigned to MP1 and MP5 modules, stratified by developmental stage (**Figure 3C**). fetal MP5 is driven by proliferative regulators (MYC, E2F1, BRCA1) ^50–52^, whereas in adults it switches to HSC-maintenance TFs (MYB, SPI1, GFI1) ^53–55^; simultaneously, adult MP1 becomes dominated by inflammatory AP-1 and NF-κB regulons (e.g., FOS, JUND, NFKB2), underscoring distinct and dynamically evolving regulatory networks of MP1 and MP5.

To identify the clinical relevance of MPs, we conducted survival analysis on the developmental program MP5. We found that an 18-gene MP5 signature was associated with poor survival in two large AML cohorts (log-rank *P* < 0.005 for both cohorts, **Figure 3D-E, Supplemental Figure 3E-F, Supplemental Table 6, Supplemental Methods**). This motivated us to create a more sophisticated model integrating multiple aging programs to quantify biological age.

### A machine learning-derived transcriptomic clock for human HSC/MPP aging

To develop a robust stem cell aging clock based on HSC/MPP transcriptomes, we expanded the analysis with our refined dataset (59,723 HSC/MPPs from 28 donors) with two independent published cohorts (17,211 HSC/MPPs from 24 donors; 4,004 HSC/MPPs from 32 donors, respectively)^16,33^, forming the largest human stem cell aging cohort comprising of 84 donors (80,938 cells). This expands cohort increased donor diversity and statistical power across the human lifespan. To reduce technical noise while preserving biological variability, we implemented a “metacell” strategy that 20 randomly sampled HSC/MPP cells from each donor were aggregated to generate one metacell. This resulted in a total of 4,088 metacells spanning ages from 4 PCW to 77 years (**Figure 4A**). Donor’s chronological age was used as the ground truth label for model training.

**Figure 4.**
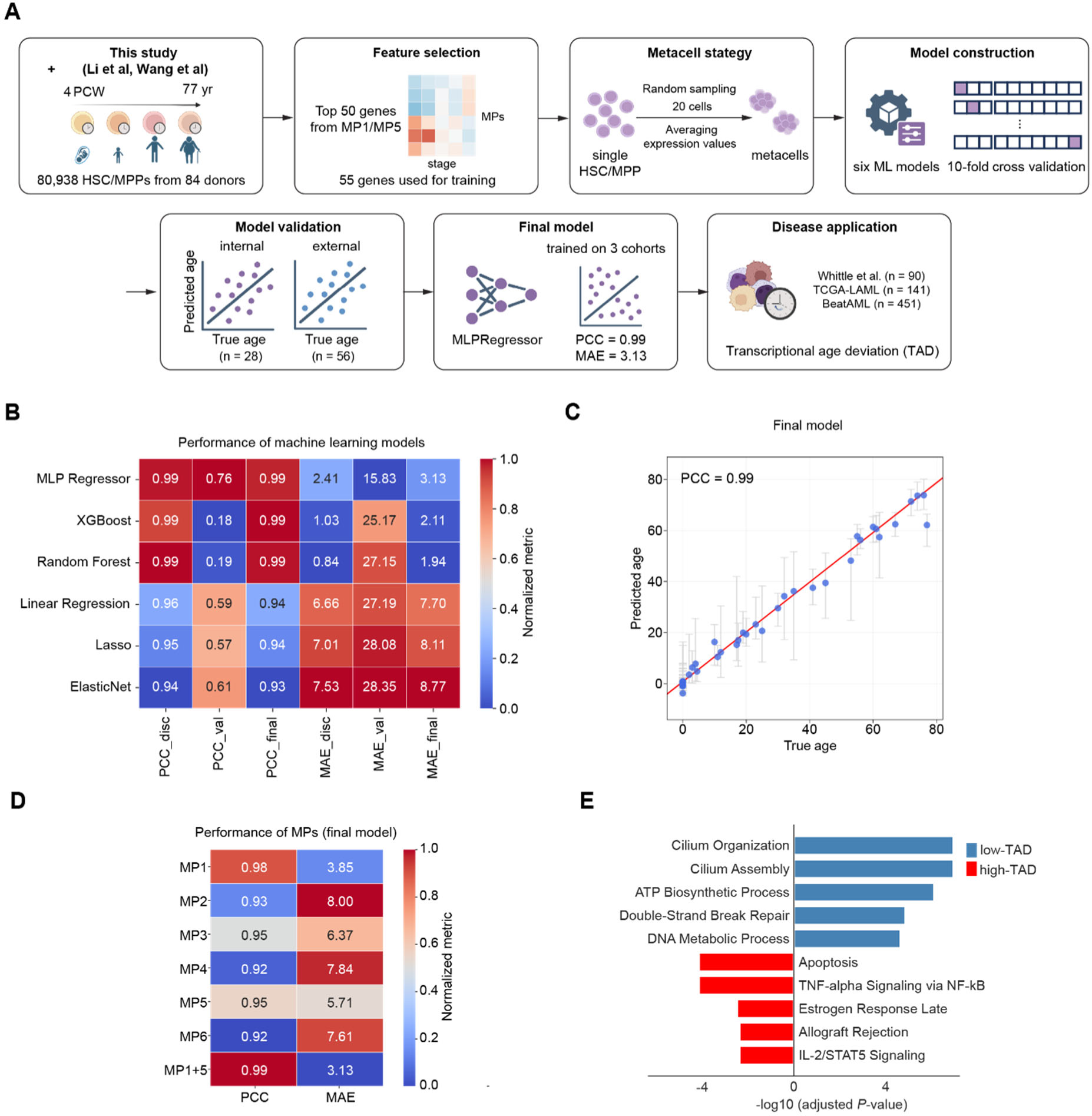
A novel HSC/MPP aging clock model across the human lifespan. (**A**) Workflow of aging clock model construction, evalutation and application. The main steps included processing of discovery and independent validation cohorts, feature selection, metacell calculation, machine learning (ML) model construction, internal and external validation, final model training on all 3 cohorts, and application of the model on AML datasets provided by Whittle et al^43^. (**B**) Performance of six common machine learning models across the discovery, validation and combined cohorts. The models are ranked by their mean absolute error (MAE) on the validation cohort. (**C**) Performance of the final model (trained on 3 cohorts). (**D**) Performance of the final model trained on 3 cohorts using each MP and a combined [MP1 + MP5] as the model’s features for training. (**E**) Pathway enrichment results using differentially expressed genes between metacells belonging to the low-TAD group and the high-TAD group.

We prioritized MP1 and MP5 for feature selection given their strongest associations with age, and extracted 55 informative genes used as input features for model construction (**Supplemental Tables 2 and 7**). We next trained and benchmarked six supervised machine learning models using a 10-fold cross-validation strategy, with our dataset as discovery dataset and the two independent cohorts as external validation datasets (**Methods**). While several models achieved similarly high Pearson correlation coefficients (PCCs) on the discovery and validation datasets (PCC = 0.99), MLPRegressor consistently demonstrated superior generalization performance, maintaining the highest PCC on the validation dataset (0.76). In contrast, XGBoost and Random Forest showed a pronounced reduction in validation PCC (≈0.18-0.19), indicating limited generalizability. Therefore, we selected MLPRegressor as our final model (**Figure 4B, Supplemental Figure 4A-B**). We retrained the final model using all three cohorts, resulting in a global stem cell aging clock with an MAE of 3.13 and a PCC of 0.99 (**Figure 4C**). To further interpret the molecular drivers underlying model predictions, we calculated permutation-importance scores for each feature in the final model, identifying genes such as *AREG, CRHBP* and *GNG11* as the most influential predictors, suggesting their relevance to age-related transcriptional reprogramming (**Supplemental Figure 4C**). Moreover, to systematically assess the contribution of each MP, we trained and compared separate models using genes derived from each MP and MP1 + MP5. This analysis confirmed that [MP1 + MP5] based models consistently outperformed those derived from other modules, highlighting the key role of these modules in defining HSC/MPP age (**Figure 4D**).

### TAD reveals biologically distinct and clinically relevant states in AML stem cells

To investigate whether hematological malignancies are associated with altered HSC/MPP biological aging, we applied the age prediction model to a large scRNA-seq cohort of acute myeloid leukemia (AML), comprising 90 patients of 73,360 single HSC/MPPs^43^ (**Supplemental Table 8**). Cells were aggregated into 3,715 HSC/MPP metacells and we calculated the transcriptional age deviation (TAD) for each metacell after applying correction to the chronological age-dependent bias (**Methods**). Metacells within the lowest and highest 10% of TAD were annotated as low-TAD and high-TAD groups, respectively (low-TAD group: TAD < −9.5, n = 334 metacells; high-TAD group: TAD > 4.8, n = 272 metacells). Differential expression analysis identified 2483 genes up-regulated in low-TAD group and 366 genes up-regulated in high-TAD group.

High-TAD group were dominated by pro-inflammatory and stress-response programs, including TNF-α signalling via NF-κB, IL-2/STAT5 signaling and apoptosis. Low-TAD cells were selectively enriched for aerobic respiration and primary cilium assembly (adjusted *P* < 0.05, **Figure 4E**). The enrichment of aerobic-respiration pathways in the low-TAD group mirrors the MP4 activity characteristic of fetal and early-postnatal HSCs. Notably, upregulated mitochondrial oxidative metabolism has been linked to the enhanced proliferative capacity of early HSCs and to drug resistance in AML^56–58^. In addition, primary cilia are reported essential for primitive hematopoiesis in both zebrafish and human and dysregulated in AML^59,60^.

Strikingly, this biologically defined TAD showed a strong correlation with established clinical risk. We observed a stepwise decrease in TAD values across European LeukemiaNet (ELN) 2022 risk groups, with adverse-risk patients exhibiting significantly lower TAD than favorable-risk patients (Kruskal–Wallis *P* = 1.4 × 10^−3^, **Supplemental Figure 4D**). This initial finding suggested that a biologically “younger” stem cell state is associated with higher-risk disease genetics.

### TAD provides independent and additive prognostic value in bulk AML cohorts

To assess the clinical utility and broad applicability of TAD, we extended our analysis to two large-scale bulk RNA-seq cohorts, TCGA-LAML (n=141) and BeatAML (n=451). Across both cohorts, TAD powerfully stratified patient survival. Patients with low TAD exhibited dramatically shorter overall survival (OS) compared to those with high TAD (TCGA-LAML: median OS 8.1 vs 15.2 months; BeatAML: median OS 7.0 vs 16.4 months; log-rank *P* < 0.001 in both cohorts; **Figure 5A-B**). Importantly, in a multivariate Cox regression model on the BeatAML cohort, after adjusting for established risk factors including age (≥60 vs <60 years) and ELN 2022 risk groups, TAD remained a highly significant, independent predictor of OS (HR = 0.95, 95% CI: 0.91-0.98, *P* < 0.005). These results firmly establish TAD as a prognostic marker with power extending beyond current clinical standards.

**Figure 5.**
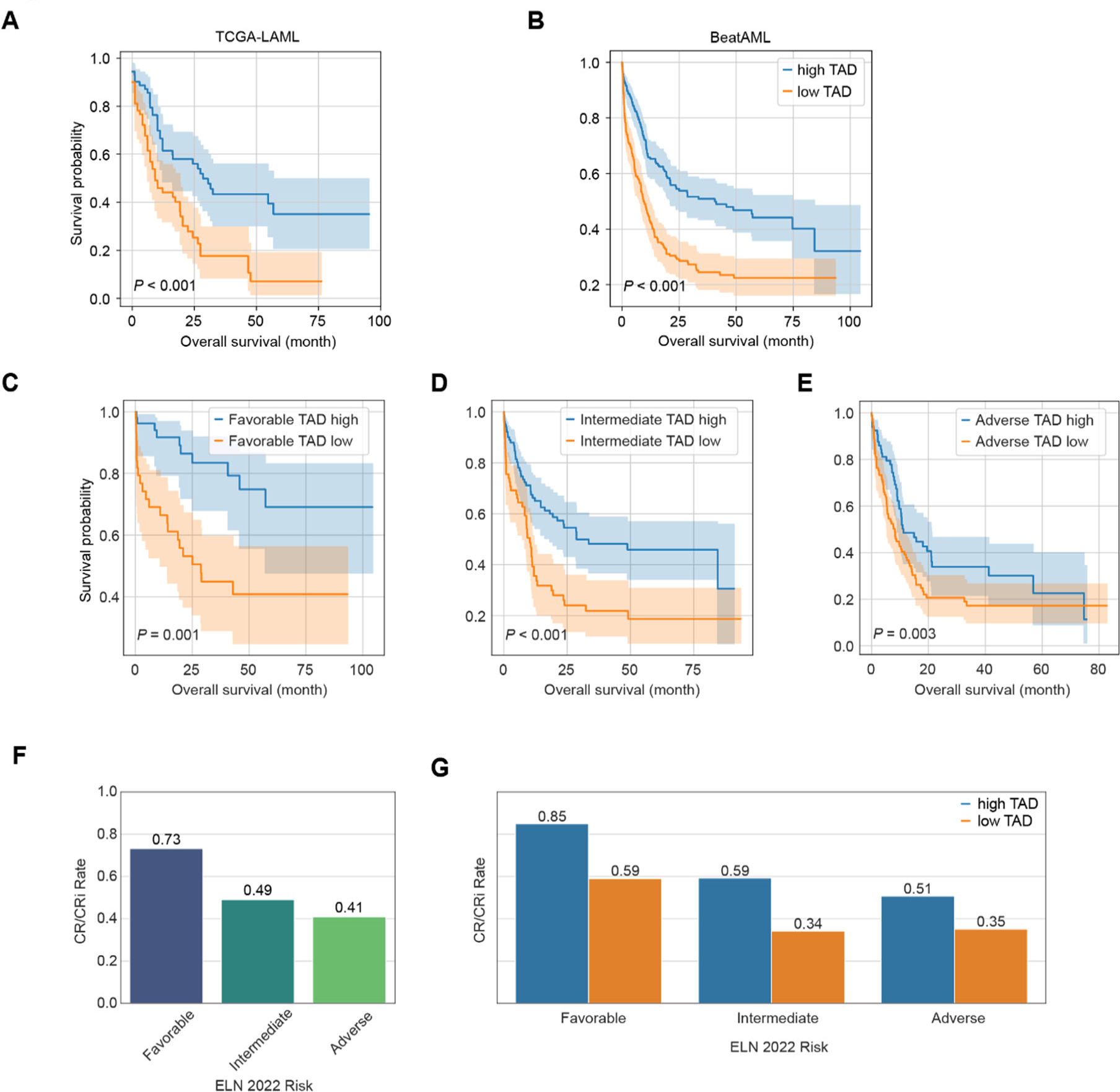
TAD stratifies risk and predicts outcome in bulk RNA-seq AML cohorts. (**A-B**) Survival analysis of AML patients divided by TAD in TCGA-LAML (**A**) and BeatAML (**B**) cohorts. (**C-E**) Survival analysis of AML patients re-stratified by TAD within the “Favorable” (**C**), “Intermediate” (**D**), and “Adverse” (**E**) ELN risk group. (**F**) Complete remission (CR/CRi) rates among patients in the three ELN risk groups. (**G**) CR/CRi rates among patients in the ELN risk groups re-stratified by TAD.

We next investigated whether TAD could resolve the known heterogeneity within ELN 2022 risk groups in the BeatAML cohort. Kaplan-Meier analysis revealed that TAD robustly re-stratified patients into distinct survival groups within each ELN category (**Figure 5C-E**). This effect was most pronounced in the clinically challenging intermediate-risk group, where patients with low TAD had a dismal median OS of just 8.6 months, compared to 18.4 months for their high-TAD counterparts (log-rank *P* < 0.001). This powerful re-stratification ability was also observed in the favorable-risk (median OS 19.6 vs 30.7 months, *P* = 0.001) and adverse-risk groups (median OS 5.4 vs 10.2 months, *P* = 0.003).

Furthermore, TAD demonstrated a significant ability to predict treatment response. While complete remission (CR) or CR with incomplete hematologic recovery (CRi) rates expectedly decreased across ELN favorable (73%), intermediate (49%), and adverse (41%) risk groups (**Figure 5F**), TAD provided additional predictive resolution. Within each ELN category, patients with low TAD consistently exhibited lower CR/CRi rates compared to their high-TAD counterparts (**Figure 5G**). This re-stratification was statistically significant in both the favorable-risk (high-TAD: 85%, low-TAD: 59%; chi-squared test, *P* = 0.009) and the clinically challenging intermediate-risk groups (high-TAD: 59%, low-TAD: 34%; *P* = 0.002). A similar trend was observed in the adverse-risk group (high-TAD: 51%, low-TAD: 35%; *P* = 0.063), collectively highlighting the potential of TAD to identify individuals unlikely to respond to standard therapy across the entire risk spectrum.

Collectively, these results establish TAD not just as a correlative biomarker, but as a quantitative metric reflecting fundamental aging-related cellular states that are co-opted in malignancy, drive clinical outcomes, and offer a valuable new dimension for risk assessment in AML.

## Discussion

In this study, we leveraged a comprehensive lifespan atlas of human hematopoiesis to develop a stem-cell-specific aging clock, leading to the definition of transcriptional age deviation (TAD), a quantitative biomarker with profound clinical implications for acute myeloid leukemia (AML). Our central finding is that TAD, which measures the degree of oncofetal reprogramming in cancer cells, serves as a powerful and independent predictor of patient survival and therapy response. We demonstrated that TAD not only stands on its own as a prognostic marker but also provides significant additive value to the current gold-standard ELN 2022 risk classification. Specifically, TAD effectively re-stratifies the clinically ambiguous intermediate-risk group, identifying a high-risk subset with dismal survival outcomes. This work bridges the fundamental biology of HSPC aging with the urgent clinical need for improved risk assessment in AML.

Our analysis provides a compelling biological explanation for the paradoxical “younger-is-worse” paradigm observed with TAD. The low-TAD state is not a sign of cellular health, but rather a hallmark of developmental reprogramming, where leukemic cells reactivate transcriptional programs characteristic of fetal hematopoiesis. These programs, particularly those related to aerobic respiration, RNA splicing, and proteostasis (as captured by our MP4 and MP5 modules), confer a highly proliferative, chemo-resistant phenotype optimized for malignant expansion. This finding aligns with the established concept that reactivation of developmental pathways is a common strategy for tumorigenesis^61–63^. Unlike traditional LSC markers (e.g., CD34, CD123) that define cellular identity^64–66^, TAD acts as a functional state descriptor, quantifying a cell’s position on a continuous aging-developmental axis and thereby offering an orthogonal layer of prognostic information.

The foundation of our clinically relevant biomarker is a meticulously constructed atlas of over 186,000 human HSPCs, spanning from early fetal development to late adulthood. This resource allowed us to delineate six robust molecular programs (MPs) governing HSPC function across the human lifespan. We identified two core, conserved aging programs—MP1 (inflammaging) and MP5 (RNA splicing and proteostasis)—whose opposing dynamics from birth to old age encapsulate the key trade-offs in HSC biology. The progressive accumulation of MP1 activity reflects the well-documented pro-inflammatory drift in the aged bone marrow niche, a known driver of myeloid skewing and clonal hematopoiesis^8–10,67^. Conversely, the sharp decline of the MP5 program after birth highlights a critical developmental window for RNA splicing and proteostasis, pathways essential for maintaining HSC fitness but which are co-opted in malignancy^11–13,68–71^. Our ability to build a highly accurate aging clock (MAE = 3.13 years) from just 55 genes derived from these two core programs underscores their central role in defining stem cell age.

Our study has several limitations. First, while our cohort of 84 donors is the largest of its kind, representation at extreme ages remains limited. Second, our analysis is based on transcriptomic data alone; future integration with epigenomic and proteomic data will provide a more comprehensive view of HSC aging and its dysregulation in disease. Third, while we validated TAD in large retrospective cohorts, its predictive utility should be prospectively evaluated in clinical trials to determine its stability and role in guiding treatment decisions.

In conclusion, our work provides a foundational map of hematopoietic aging and establishes TAD as a robust, clinically translatable biomarker. By quantifying the deviation from a healthy aging trajectory, we have created a powerful new tool for precision oncology that converts a complex biological signature into a single, interpretable metric. The strong performance of TAD in bulk RNA-seq cohorts highlights its potential for immediate implementation in clinical settings. This study demonstrates that a deep understanding of fundamental aging biology can unlock novel strategies for stratifying disease risk and personalizing therapy in AML and potentially other hematologic malignancies.

## Authorship

Contribution: Conceptualization: G.W. & H.C.; Methodology: G.W. & H.C.; Formal analysis: H.C. & P.D.; Investigation: H.C. & J.X.; Data curation: H.C.; Writing: G.W. & H.C.; Visualization: H.C.; Supervision: G.W.; Funding acquisition: G.W. Conflict of interest disclosure: All authors declare no conflict of interest.

